# Human-specific paralogs of SRGAP2 induce neotenic features of microglia structural and functional maturation

**DOI:** 10.1101/2024.06.28.601266

**Authors:** Carlos Diaz-Salazar, Marine Krzisch, Juyoun Yoo, Patricia R. Nano, Aparna Bhaduri, Rudolf Jaenisch, Franck Polleux

## Abstract

Microglia play key roles in shaping synaptic connectivity during neural circuits development. Whether microglia display human-specific features of structural and functional maturation is currently unknown. We show that the ancestral gene *SRGAP2A* and its human-specific (HS) paralogs *SRGAP2B/C* are not only expressed in cortical neurons but are the only HS gene duplications expressed in human microglia. Here, using combination of xenotransplantation of human induced pluripotent stem cell (hiPSC)-derived microglia and mouse genetic models, we demonstrate that (1) HS SRGAP2B/C are necessary and sufficient to induce neotenic features of microglia structural and functional maturation in a cell-autonomous manner, and (2) induction of SRGAP2-dependent neotenic features of microglia maturation non-cell autonomously impacts synaptic development in cortical pyramidal neurons. Our results reveal that, during human brain evolution, human-specific genes SRGAP2B/C coordinated the emergence of neotenic features of synaptic development by acting as genetic modifiers of both neurons and microglia.

## INTRODUCTION

Microglia play key roles during brain development, neural circuit homeostasis, and pathology ^1–3^. As the resident immune cells of the central nervous system (CNS), they provide essential support for proper neuronal development, synaptic maturation, and function, both embryonically and postnatally ^1,2^. In mammals, from rodents to primates including humans, adult neuronal connectivity is reached during postnatal development through a combination of synapse formation, maturation and elimination ^4–6^. Microglia are critical for shaping and maintaining synaptic connectivity in various circuits by regulating synapse development, maintenance and function ^1,2,7–9^. Microglia can also respond to and modulate neuronal activity at the synaptic level ^10,11^. Additionally, as a tissue-resident macrophage, they continually survey their environment, searching for and quickly responding to injury cues, and phagocytosing any debris, pathogens, or injured cells they may encounter ^9,12^.

Microglia originate from the embryonic yolk sac and migrate to the brain early during development (around Embryonic Day (E)8.5 in mice and gestational week (GW) 5 in humans), where they mature *in situ* ^13–15^. However, like neurons, microglia development continues postnatally ^1,2^. In mice, microglia transition from an ameboid shape into a highly complex and branched morphology, while acquiring expression of a set of genes allowing them to sense their environment and perform their surveillance and phagocytic capabilities ^16–18^. Importantly, in mice, the timing of postnatal microglial structural and functional maturation largely coincides with that of postnatal synaptic maturation, highlighting the importance of synchronizing both cellular processes at the molecular level ^1,6^.

While the dynamics and timing of neuronal and microglial maturation have been well-characterized in the brains of rodents ^1,2,6,15,19–22^ and to some extent in non-human primates ^23,24^, less is known about the timing and features of microglia maturation in the human brain ^4,15,25^. In brain structures like the neocortex, one the most salient and striking feature characterizing human neuron development is the slow, neotenic, timing of their morphological and functional maturation and, in particular, synaptic maturation taking weeks to months in most mammals from rodents to non-human primates versus decades after birth in humans ^26–29^. This strikingly prolonged period of synaptic development and maturation is thought to extend critical periods of circuit remodeling enabling longer periods of postnatal development when circuits can be modified by experience-dependent plasticity ^30^.

Whether microglia development and maturation exhibit human-specific features is still largely unknown. While recent attempts characterizing the transcriptional state of fetal and postnatal human microglia have pointed to clear differences between these two stages, the precise timing and magnitude of such changes are still poorly characterized, impairing our ability to identify candidate molecular effectors of microglia structural and functional maturation during development and determine if some of these effectors act as species-specific genetic modifiers of microglia development and function ^31–34^. Current efforts establishing native-like models to study human microglia function, such as those using microglia derived from human induced pluripotent stem cells (hiPSC), have been largely focused on proof-of-concept modeling and applicability to disease models where microglia play an important role, and for example highlighted certain human-specific features of microglial response to injury or pathological states such as neurodegeneration ^35,36^. Here, we provide evidence that human microglia exhibit human-specific morphological and transcriptomic features, characterized by a slow maturation trajectory. We characterize the role of one genetic driver of such features, the ancestral gene SRGAP2A, acts as a master regulator of microglial maturation and function. We demonstrate that human-specific paralogs SRGAP2B/C, through inhibition of SRGAP2A, induce neotenic features of microglia morphological and functional maturation which in turns affects synaptic development in neurons in a cell non-autonomous manner.

## RESULTS

### Human microglia show features of neotenic development

We first assessed the timing of morphological maturation of endogenous microglia in the mouse neocortex, both in terms of cell number and density and morphological complexity using Iba1 as a marker ^37^. Consistent with previous reports, the number of Iba1+ microglia in the mouse cortex rapidly increases during the first two weeks after birth, then declines and stabilizes around the fourth postnatal (PN) week (**Fig. S1A-B**) ^38^. We then quantified microglial morphological complexity using Sholl analysis (**Fig. S1C-E**), and, consistent with the literature ^16^, we observed that microglial branching complexity increases linearly during the first 2 postnatal weeks of development (**Fig. 1A-B**). Interestingly, we also found that following this period of hyper-ramification, microglia complexity is reduced and reach adult levels approximately 1 month after birth (**Fig. 1A-B**).

**Figure 1.**
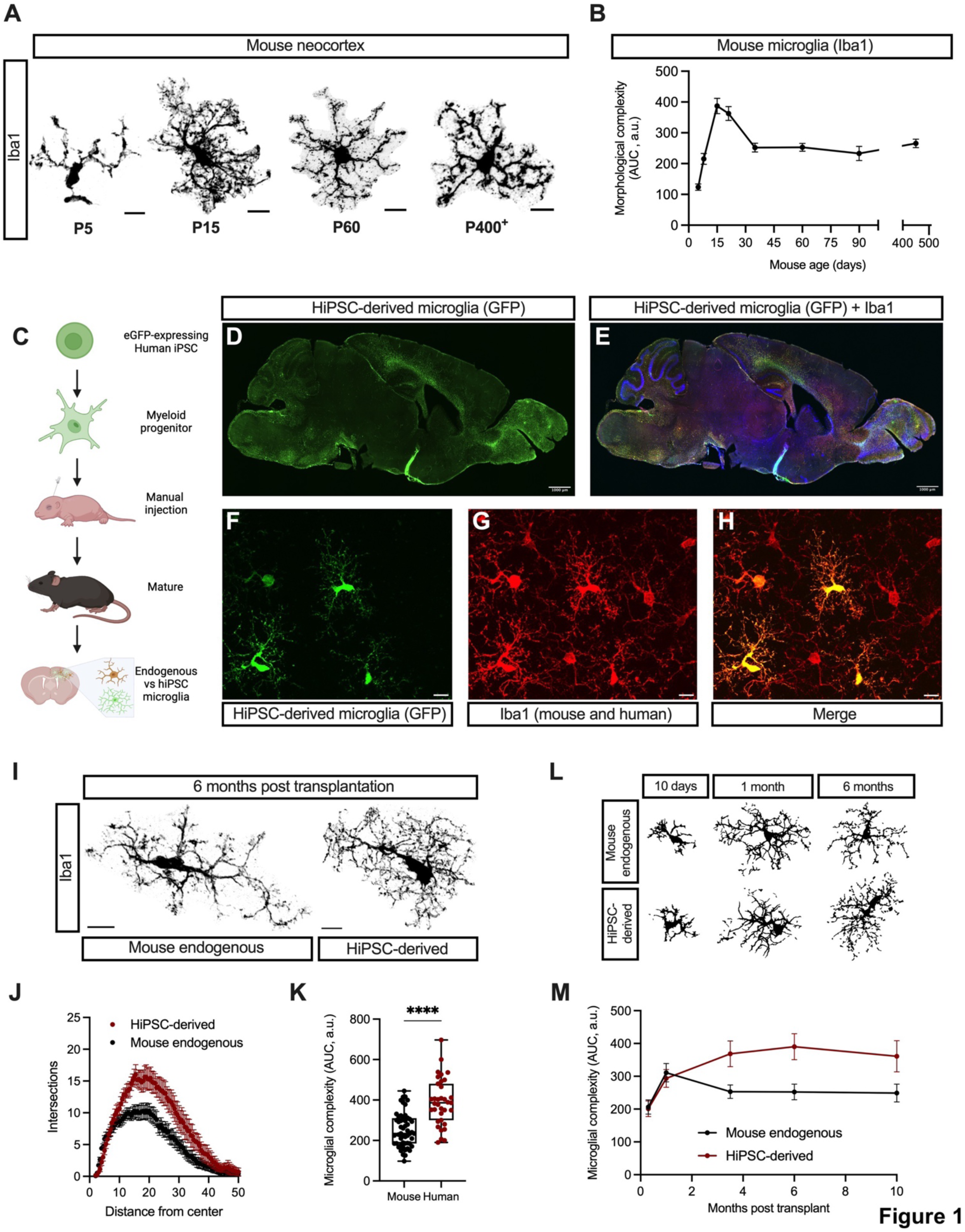
Human iPSC-derived microglia exhibit increased morphological complexity and delayed morphological maturation compared to mouse microglia. (A) Representative images of microglia from early postnatal (P5), juvenile (P15), adult (P60), and aged (P400+) mouse brains, as revealed by Iba1 staining. Scale bar, 10 μm. (B) Quantification microglial morphological complexity over time, computed as the area under the curve (AUC) of each microglia’s sholl traces. Data from three to eight mice per timepoint, 9-20 microglia per mouse. (C) Experimental approach to comparatively assess human microglial morphology. Human induced pluripotent stem cells (iPSCs) stably expressing GFP are differentiated into myeloid precursors, xenotransplanted in immunodeficient neonatal (P1-3) mice, and left to differentiate into microglia and mature until mouse adulthood. (D-H) Representative images of a 100 µm-thick sagittal section of a mouse brain xenotransplanted with GFP+ human-iPSC derived (hMG) microglia (D) and stained with Iba1 (E). Confocal maximum intensity projections revealing the morphology of hMG (F) and mouse endogenous microglia (G) on the same field of view (H) at high magnification. Scale bar, 10 μm. (I) Representative images of hMG and mouse endogenous microglia from a 6 month-old mouse. Scale bar, 10 μm, Comparison of sholl analysis (J) and morphological complexity (computed as the AUC of sholl traces) (K) of hMG and mouse endogenous microglia. Data from three mice, 8-22 cells per mouse. (L) Representative traces of hMG and mouse microglia at different developmental stages. (M) Morphological complexity of mouse and hMG microglia across time. Data from three to six mice per timepoint, 7-22 cells per mouse.

To assess morphological maturation of human microglia *in vivo*, we xenotransplanted hiPSC-derived microglia into mouse brains, and compared their morphological state to the neighboring mouse endogenous microglia (**Fig. 1C**). In this system, hiPSCs are differentiated into myeloid precursors which, upon xenotransplantation into neonatal mouse brains from immunodeficient Rag2 knockout mice engraft extensively, differentiate into microglia, and closely recapitulate the transcriptional signature of primary human microglia at diverse developmental stages (**Fig. 1D-H**) as previously shown ^31,36,39,40^. At 6 months post-transplantation (MPT), xenotransplanted hiPSCs-derived microglia (hMG) were significantly more ramified than their mouse endogenous counterparts (**Fig. 1I-K**). We then assessed their hMG morphological maturation longitudinally and found a marked delay in reaching peak morphological complexity and failure to adopt a simplified mature morphology even at 10 MPT, suggesting that their morphological maturation is neotenic under these conditions (**Fig. 1L-M**).

Microglial maturation states can also be inferred from their transcriptional signatures. In the mouse, it is widely accepted that microglia adopt a mature transcriptional signature at 2 weeks after birth ^19,22^. However, despite studies highlighting the differences between fetal and postnatal microglia, a comprehensive transcriptional signature of human microglia maturation is not yet available ^31,32^. To generate a transcriptional atlas of human microglia maturation, we used a publicly available single-nucleus RNAseq dataset containing primary human microglia and extracted their transcriptional trajectories from the second gestational trimester to adulthood ^41^. Weighted gene correlation network analysis (WGCNA) identified 166 modules of developmentally co-expressed genes, which we then further grouped into 5 highly correlated clusters of modules using a k-means unsupervised clustering algorithm (**Fig. S2A-B**). Most gene expression modules showed limited developmental expression dynamics and clustered tightly together, suggesting that the expression of the genes belonging to such modules remains stable throughout microglial maturation (**Fig. S2A-B**). However, certain modules showed a marked change in module activity throughout early human development and did not plateau until approximately 5 years of age. To confirm that this approach can infer absolute-time developmental trajectories, we applied the same methodology to a publicly available single-cell RNAseq dataset of mouse microglia ^42^, and identified similar trajectories to those observed in human microglia (**Fig. S2C-D**). Mainly, the activity of most modules remained flat during development, and those that showed a developmental change in activity reached activity plateau at 2 weeks after birth, consistent with previous reports ^22^. This data indicates that mouse and human microglia have similar developmental trajectories, but that the absolute timing of human microglia maturation is profoundly delayed compared to mouse microglia.

Of the temporally regulated gene modules in human microglia, module 160 showed the steepest increase in gene expression module activity throughout human development, reaching peak activity at around 5-10 years of age (**Fig. 2A**). In accordance with our previous findings, the activity of the mouse orthologs of the genes contained in module 160 peaked at 2-3 weeks PN (**Fig. 2B**), highlighting that human and mouse microglia share a common developmental profile albeit at very different time scales, with clear signature of transcriptional neoteny for human cortical microglia.

**Figure 2.**
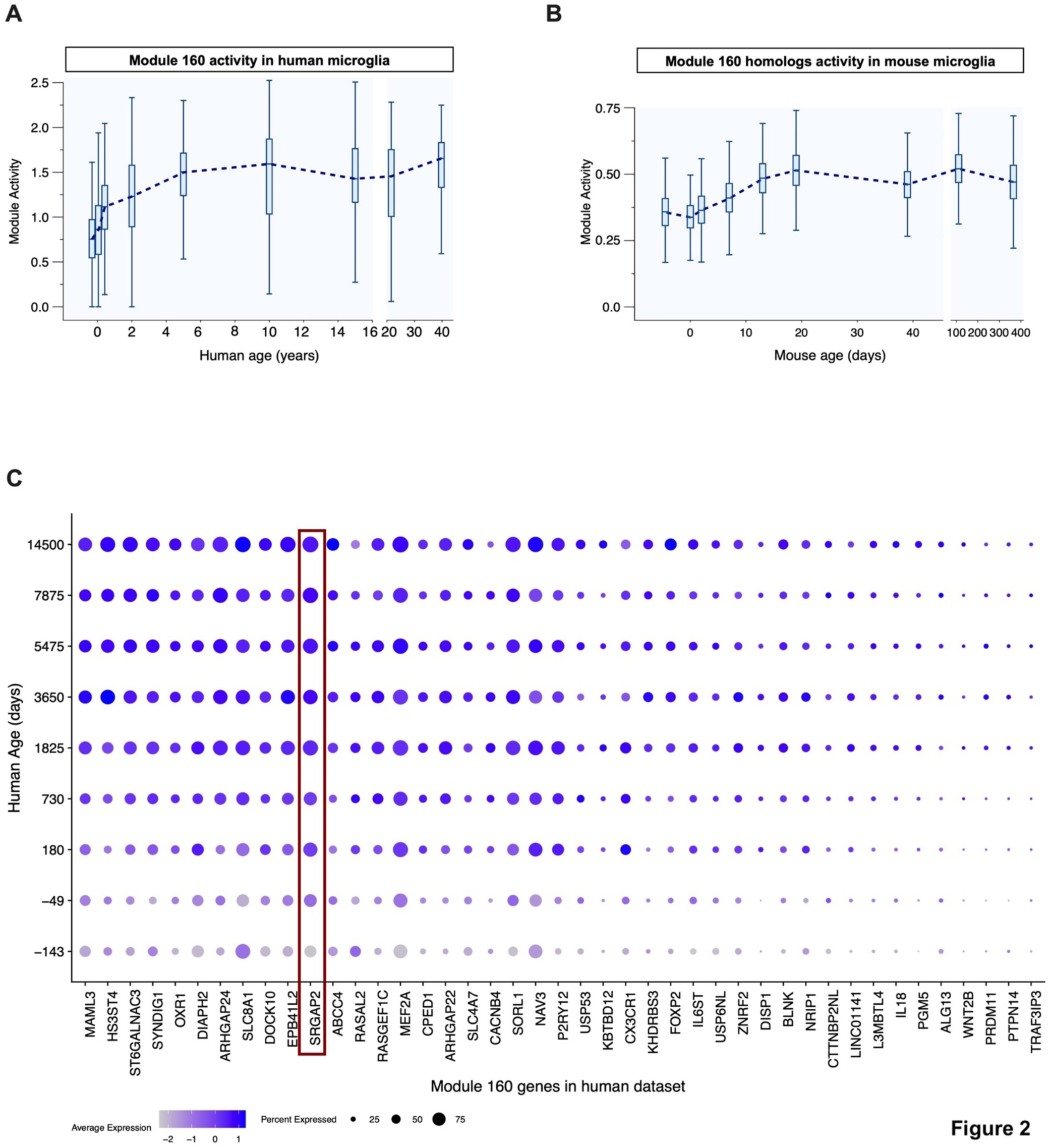
Human microglia display transcriptomic features of neotenic development. (A) Average activity throughout human development of genes contained in human module 160. Dashed line connects the median activity for each timepoint. (B) Average activity throughout mouse development of the mouse homologs to the genes contained in human module 160. (C) Dotplot heatmap representing developmental expression of the 43 human genes contained in module 160. Note the Human-specific duplicated gene Srgap2 boxed in red.

Interestingly, module 160 contains genes essential for microglial homeostasis such as *cx3cr1* and *p2ry12*, transcription factors and coactivators, and genes putatively involved in regulating cytoskeletal dynamics by modulating the activation state of small Rho-GTPases (**Fig. 2C**). Furthermore, genes in this module 160 are highly overrepresented in a primary human microglia-specific gene signature that hMG models cannot fully recapitulate ^36^, suggesting that peak expression of module 160 genes is necessary for the complete terminal maturation of human microglia.

### Ancestral SRGAP2 regulates structural maturation of microglia

To identify potential candidate genes acting as human-specific modifiers of microglia morphological and/or functional maturation, we examined human-specific gene duplications. We and others recently demonstrated that human-specific gene duplications represent a potent source of human-specific genomic modifiers of neuronal development and connectivity, ultimately impacting cortical circuit function ^30,43–47^. Interestingly, module 160 contains SLIT-ROBO Rho GTPase activating protein 2 (*Srgap2* in mammals, named *SRGAP2A* in human), a gene that has undergone two main partial duplications leading to the emergence of the human-specific paralogs *Srgap2b* and *Srgap2c* approximately 3.5-2.4 million years ago (mya) ^44,48^. The ancestral gene Srgap2 (present in all mammals) and its human-specific paralogs *Srgap2b/c* are the only one of ∼31 cognate HSGD (red stars in **Fig. 3A**) strongly expressed in human microglia. Furthermore, *Srgap2* temporal expression pattern in mouse and *Srgap2a* and its human-specific paralogs *Srgap2b/c* in human microglia coincides with terminal microglia maturation (**Fig. S2E-F**), suggesting that *Srgap2b/c* are potentially unique candidates for controlling human-specific features of microglial morphological and/or functional maturation.

**Figure 3.**
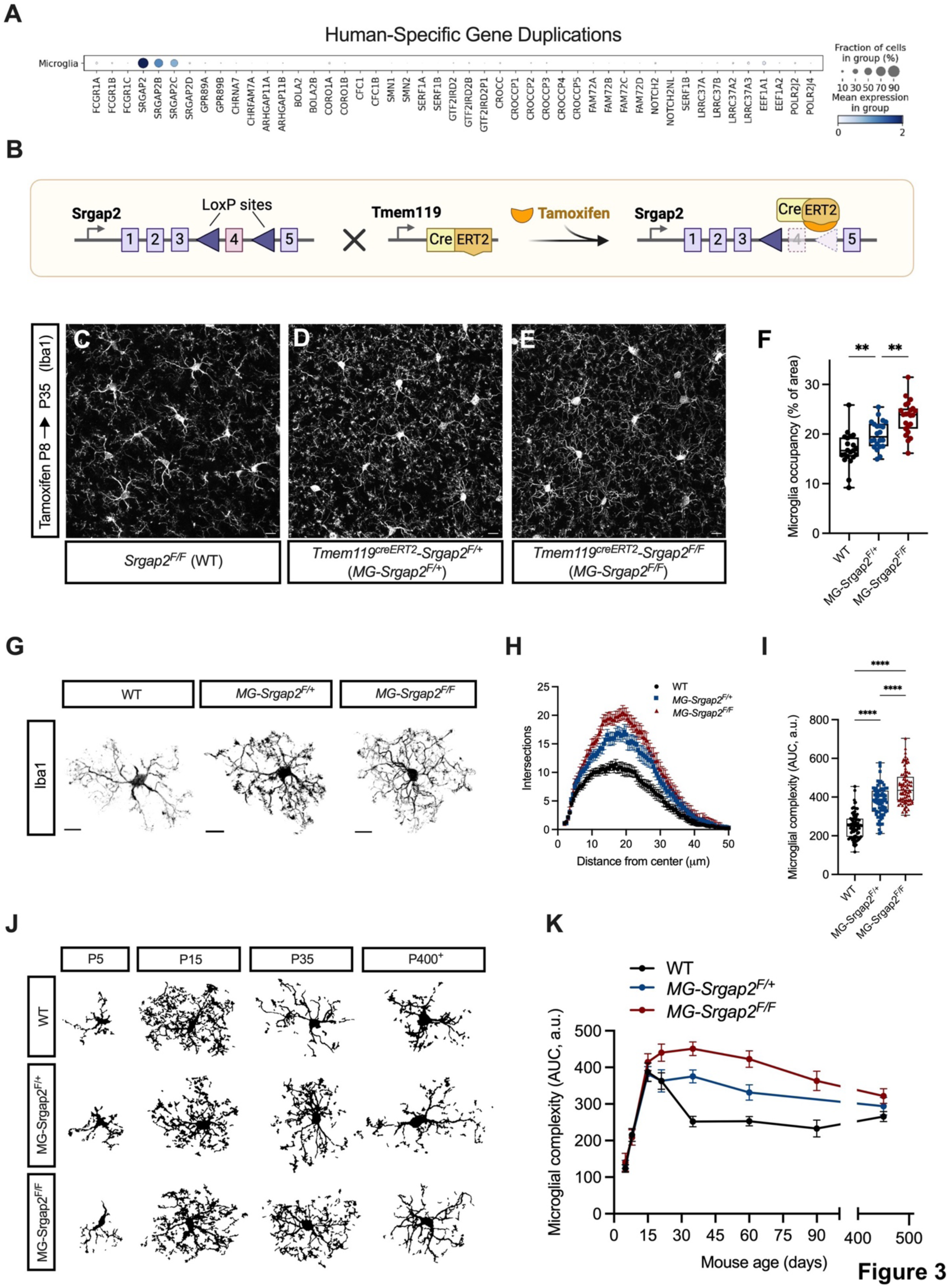
Srgap2 is a master regulator of microglial maturation. (A) Dotplot heatmap depicting the expression of all 31 annotated human-specific gene duplications families (red stars) and their cognate ancestral gene copy in primary human microglia. (B) Schematic representation of the genetic approach used to conditionally ablate Srgap2 in microglia. C-E) Representative images of microglia (as revealed by Iba1 staining) from WT (*Srgap2^F/F^*), *MG-Srgap2^F/+^* (*Tmem119^CreErt^*^2^ *Srgap2^F/+^*), and *MG-Srgap2^F/F^* (*Tmem119^CreErt^*^2^ *Srgap2^F/F^*) P35 mice. Scale bar, 10μm. (F) Quantification of area covered by microglia per field of view from these mice. Data from eight mice per genotype, two to four fields per mouse. (G) Representative images of individual microglia from these mice. Scale bar, 10μm. H and I) Comparison of sholl analysis (H) and morphological complexity (I) of microglia from these mice. Data from six mice per genotype, 11-17 cells per mouse. (J) Representative traces of microglia from these mice at different ages. (K) Morphological complexity of microglia from these mice across time. Data from three to eight mice per timepoint and genotype, 8-22 cells per mouse.

In human neurons, *Srgap2b/c* encode two almost identical (only 6 non-synonymous base pair mutations), truncated and intrinsically unstable proteins that remain able to interact with the ancestral full-length SRGAP2A protein and target it for proteosome degradation, inhibiting all known functions of SRGAP2A in cortical pyramidal neurons (CPNs) and phenocopying SRGAP2 haploinsufficiency (**Fig. S2G**) ^44,45,49,50^. We therefore hypothesized that expression of SRGAP2B/C in human microglia should cell-autonomously phenocopy a partial SRGAP2A loss-of-function and might modulate microglia morphological and functional maturation in a human-specific manner.

We first tested the phenotypic consequence of SRGAP2 loss of function in mice by generating a microglia-specific *Srgap2* deletion by crossing a microglia-specific and inducible *Tmem119-Cre^ERT^*^2^ with a conditional knockout mouse model (*Srgap2^F/F^*) (**Fig. 3B**) ^51,52^. Mice were treated with tamoxifen early during development (P8-9), achieving high recombination efficiency in cortical microglia (**Fig. S3A-E**), to drive Srgap2 deletion. Deletion of 1 copy (*MG-Srgap2^F/+^*) or 2 copies (*MG-Srgap2^F/F^*) of *Srgap2* specifically in microglia led to a dose-dependent increase in microglia tissue coverage of the mouse cortex, when compared to littermate, tamoxifen-treated control mice (WT i.e. *Srgap2 ^F/F^*) (**Fig. 3C-F**). However, adult cortical microglia cell density was unchanged in mice from all genotypes (**Fig. S3F-I**), suggesting that the increase coverage may be due to increased microglia morphological complexity. Indeed, *Srgap2*-deficient microglia showed a significant, dose-dependent, increase in complexity, displaying a hyper-ramified morphology at P35 (**Fig. 3G-I**). Notably, microglia from constitutive Srgap2 heterozygous knockout mice (*Srgap2^+/-^*) similarly displayed a hyper-ramified phenotype compared to control wild-type littermates at P35 (**Fig. S3J-L**).

Since *Srgap2* expression monotonically increases in mouse cortical microglia during the first 3 postnatal weeks of development and is maintained throughout adulthood (**Fig. S2E**), we determined when this increase morphological complexity emerged during postnatal development. After being treated with 4-hydroxy-Tamoxifen early after birth (P2-3), both *MG-Srgap2^F/+^* and *MG-Srgap2^F/F^* microglia reached peak morphological complexity at 2 to 4 weeks after birth, but then display a delayed period of morphological simplification, only approaching WT morphological complexity levels after one year of age (**Fig. 3J-K**). Altogether, our data suggests that SRGAP2 negatively regulates microglial morphological complexity and that its deletion results in a dose-dependent delay in morphological maturation i.e. neotenic retention of ‘immature’, hyper-ramified, morphology.

We then tested whether we could capture a transcriptional signature in Srgap2-deficient microglia consistent with that of a less mature, neotenic state. Bulk RNA sequencing of FACS-sorted wild-type and *MG-Srgap2^F/F^*microglia from P30 mice, following tamoxifen induction at P8-9 (**Fig. S4A**), revealed significantly higher expression of transcripts strikingly enriched for genes encoding cytoskeletal organization and actin dynamics, such as *Tmsb4x*, *Lsp1*, and *Stmn1*, compared to WT isochronic microglia (**Fig. S4B**). Indeed, Gene Ontology enrichment analysis of the top 100 overexpressed genes in *MG-Srgap2^F/F^* microglia revealed a marked enrichment in biological processes involved in actin filament polymerization and regulation of cytoskeleton organization (**Fig. S4C**). Interestingly, the expression pattern of these genes with increased expression in *MG-Srgap2^F/F^* microglia compared to WT control microglia was highly anti-correlated with microglia maturation (**Fig. S4D**), suggesting that the transcriptional signature of Srgap2-deficient microglia is more similar to transcriptional signature found in WT mice at earlier stages of development. In line with our previous findings, this data supports our morphological observation that SRGAP2 expression is necessary for terminal morphological maturation of cortical microglia.

We finally investigated whether the neotenic maturation of *MG-Srgap2^F/F^*microglia had an impact in their functional properties. We first assessed the homeostatic surveillance capacity of *MG-Srgap2^F/F^*microglia and its WT counterparts using *in vivo* 2-photon (2P) microscopy (**Fig. 4A**), for which each strain was additionally crossed with a Cre-dependent tdTomato red fluorescent reporter (*Ai9* mouse line) ^53^. Semiautomatic analysis of microglia process dynamics using the MotiQ pipeline ^54^ revealed that Srgap2-deficient microglia scanned a larger area over a 10-minute time period (**Fig. 4B-C**) and featured more motile processes than WT microglia even when normalized to total area occupied by microglia processes (**Fig. 4D**). Finally, we also observed a decrease in the microglial area covered by the late phagolysosomal marker CD68, suggesting that *MG-Srgap2^F/F^* microglia have a reduced phagocytic capacity (**Fig. 4E-G**). In summary, our data suggests that *Srgap2* is necessary for terminal mouse microglia maturation, acting as a master regulator of microglial morphology and function.

**Figure 4.**
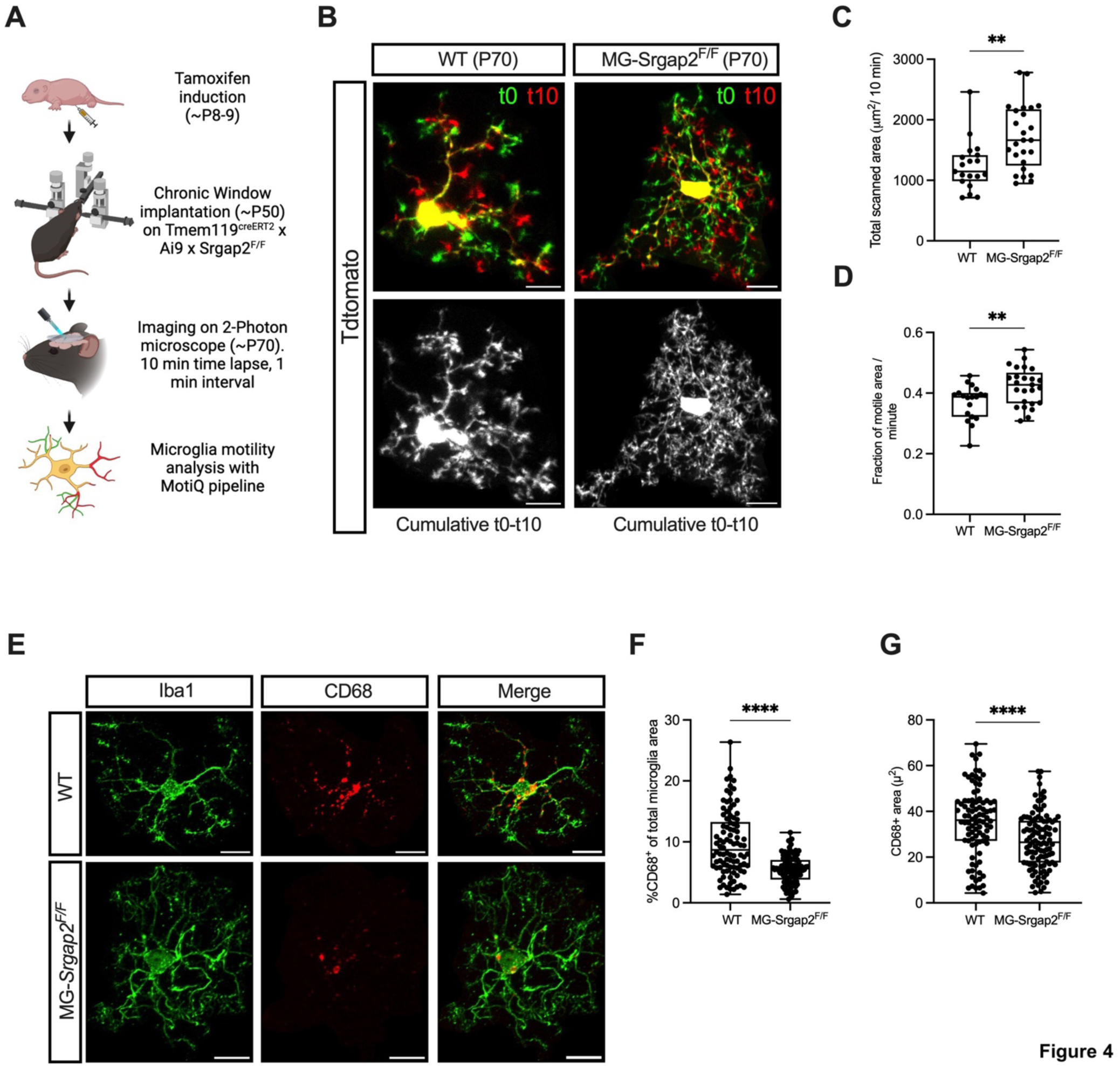
Srgap2 regulates microglial surveillance and phagocytic potential. (A) Experimental approach to assess microglial surveillance *in vivo*. (B) Representative still frames (top) and cumulative scanning (bottom) of microglia from WT and *MG-Srgap2^F/F^* mice expressing the fluorescent reporter Tdtomato, imaged over a 10-minute period. Scale bar, 10μm (C) Comparison of the total area scanned by WT and *MG-Srgap2^F/F^*microglia over a 10-minute period. (D) Average microglia motility from WT and *MG-Srgap2^F/F^* mice. Data from three mice per genotype, 19-25 cells per genotype. (E) Representative images of Iba1^+^ microglia from WT and *MG-Srgap2^F/F^* mice co-stained with the phago-lysosomal marker CD68. Scale bar, 10μm. (F) Quantification of the percentage of CD68^+^ area per Iba1^+^ microglial area from WT and *MG-Srgap2^F/F^* microglia. (G) Quantification of total CD68^+^ area in WT and *MG-Srgap2^F/F^*microglia. Data from five mice per genotype, 10-29 cells per mouse.

### Expression of human-specific Srgap2c paralogs in mouse microglia is sufficient to induce increased morphological complexity

We next tested if, as previously shown in cortical pyramidal neurons, ^44,45,49^ expression of human-specific *Srgap2c* phenocopies Srgap2 loss-of-function in microglia. We used a previously characterized transgenic mouse model ^46,51^, where we knocked-in a LoxP-STOP-LoxP:SRGAP2C expression cassette in the Rosa26 locus (*Srgap2c^LSL^*), enabling expression of SRGAPC in a Cre-dependent way. We generated crosses between *Tmem119-cre^ERT^*^2^ mouse line and *Srgap2^LSL^* mice and induced Cre-dependent Srgap2C expression following tamoxifen induction at P8-9 (**Fig. 5A**) ^46^. Analysis at P40 revealed that mouse Iba1+ microglia expressing the human SRGAP2C exhibit increased morphological complexity (**Fig. 5B-D**), to levels similar to Srgap2-deficient microglia at the same age (**Fig. 3B-F**).

**Figure 5.**
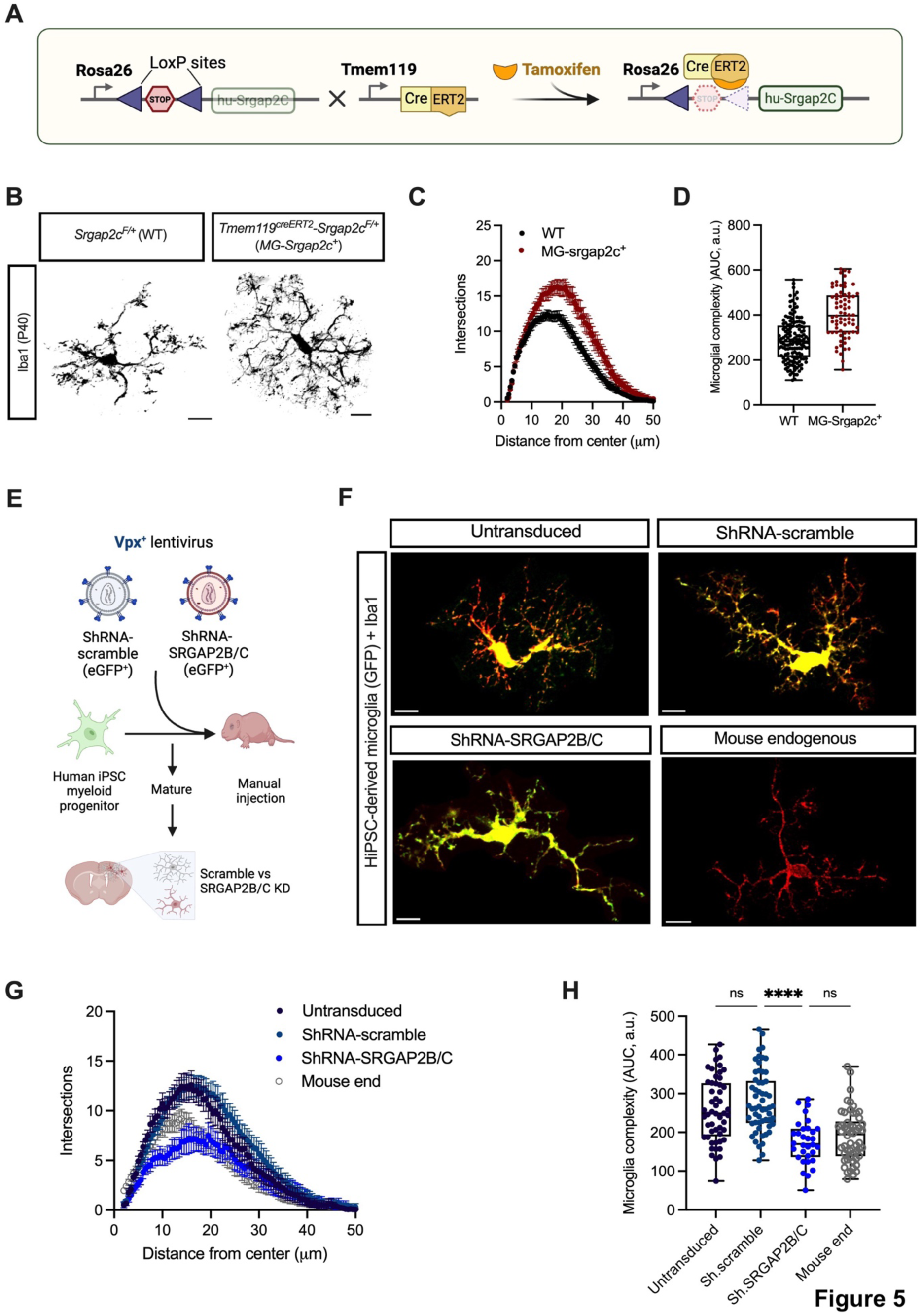
The human-specific *Srgap2c* gene modulates microglial morphology. (A) Schematic representation of the genetic approach used to ectopically express the human gene Srgap2c in mouse microglia. (B) Representative images of individual microglia from WT and *MG-Sgrap2c^+^*(*Tmem119^CreErt^*^2^ *Srgap2c^F/+^*) mice. Scale bar, 10μm. (C-D) Comparison of sholl analysis (C) and morphological complexity (D) of microglia from these mice. Data from four to eight mice per genotype, 14-26 cells per mouse. (E) Experimental strategy to knockdown human-specific *Srgap2* paralogs in hMGs. Human iPSC-derived myeloid progenitors were transduced with a EGFP-expressing lentivirus also encoding short-hairpin (sh) RNA targeting the human-specific paralogs *Srgap2b/c*, scrambled shRNA, or untransduced. (F) Representative images of untransduced hMGs, lentivirally-transduced hMGs, or control endogenous microglia from 3-month-old mice. Scale bar, 10μm. (G-H) Comparison of sholl analysis (G) and morphological complexity (H) of microglia from these mice. Data from three mice per condition, 32-58 cells per condition.

### Deletion of human-specific Srgap2b/c paralogs in human microglia leads to reduction morphological complexity

We then assessed whether the human-specific genes *Srgap2b/c* were necessary to maintain the high morphological complexity observed in human microglia. To do so, we used modified lentiviral particles packaging the Simian Immunodeficiency Virus Vpx protein on their capsid to infect human iPSC-derived myeloid precursors. The Vpx protein has been shown to promote lentiviral infection of myeloid cells, and to greatly increase transduction efficiency ^55,56^. These lentiviral particles contained either control (Scrambled) short-hairpin RNA (shRNAs) or shRNA targeting the shared 3’UTR region of the human-specific paralogs *Srgap2b and Srgap2c*, which effectively downregulates the expression of both human-specific paralogs as previously characterized ^44,50^. Following lentiviral infection of human iPSC-myeloid progenitors, we performed their xenotransplantation into the cortex of *Rag2^-/-^* knockout mouse neonates, where they differentiated into human microglia-like cells (**Fig. 5E**). We found that the morphological complexity of control cells transduced with control scrambled shRNA lentivirus was similar to that of untransduced hMG at 3 months post-transplant (**Fig. 5F-H**). Strikingly, cells transduced with shRNA *Srgap2b/c* lentivirus displayed a significant decrease in morphological complexity compared to untransduced human microglia or human microglia expressing control scrambled shRNA at 3 MPT (**Fig. 5F-H**). This result demonstrate that human-specific paralogs *Srgap2b/c* are required for the morphological complexity observed in human microglia *in vivo*.

### Microglia-specific Srgap2 deletion impacts synaptic development cell non-autonomously in the mouse cortex

Microglial cells are required for proper synaptic development and have been involved in synapse formation and elimination ^1,2,7,8^. Given that in the mouse, synaptic formation and maturation peaks during the second and third postnatal weeks, corresponding with the peak of microglial morphological complexity, we tested if the delayed microglial maturation observed in *MG-Srgap2^F/F^* mice could affect the timing of synaptic development in cortical pyramidal neurons non cell-autonomously. To assess synaptic development and structural dynamics, we used *in utero* electroporation (IUE) to sparsely label layer 2/3 Pyramidal Neurons (L2/3 PNs) in the somatosensory cortex, using a FlpO-recombinase dependent expression system (**Fig. 6A**). Briefly, *MG-Srgap2^F/F^* or control WT littermate mouse embryos were electroporated with plasmids encoding red fluorescent protein mScarlet flanked by FRT regions, and low amounts of a plasmid encoding the recombinase FlpO enabling sparse recombination and expression of mScarlet in L2/3 PNs (**Fig. 6B**). This manipulation enables visualization of dendritic spines in L2/3 PNs in wild-type mice or mice where Srgap2 is deleted in nearly all microglia (*MG-Srgap2^F/F^*). We verified that the tamoxifen induction in these mice resulted in increased morphological complexity of Iba1+ microglia compared to control mice (**Fig. 6C**).

**Figure 6.**
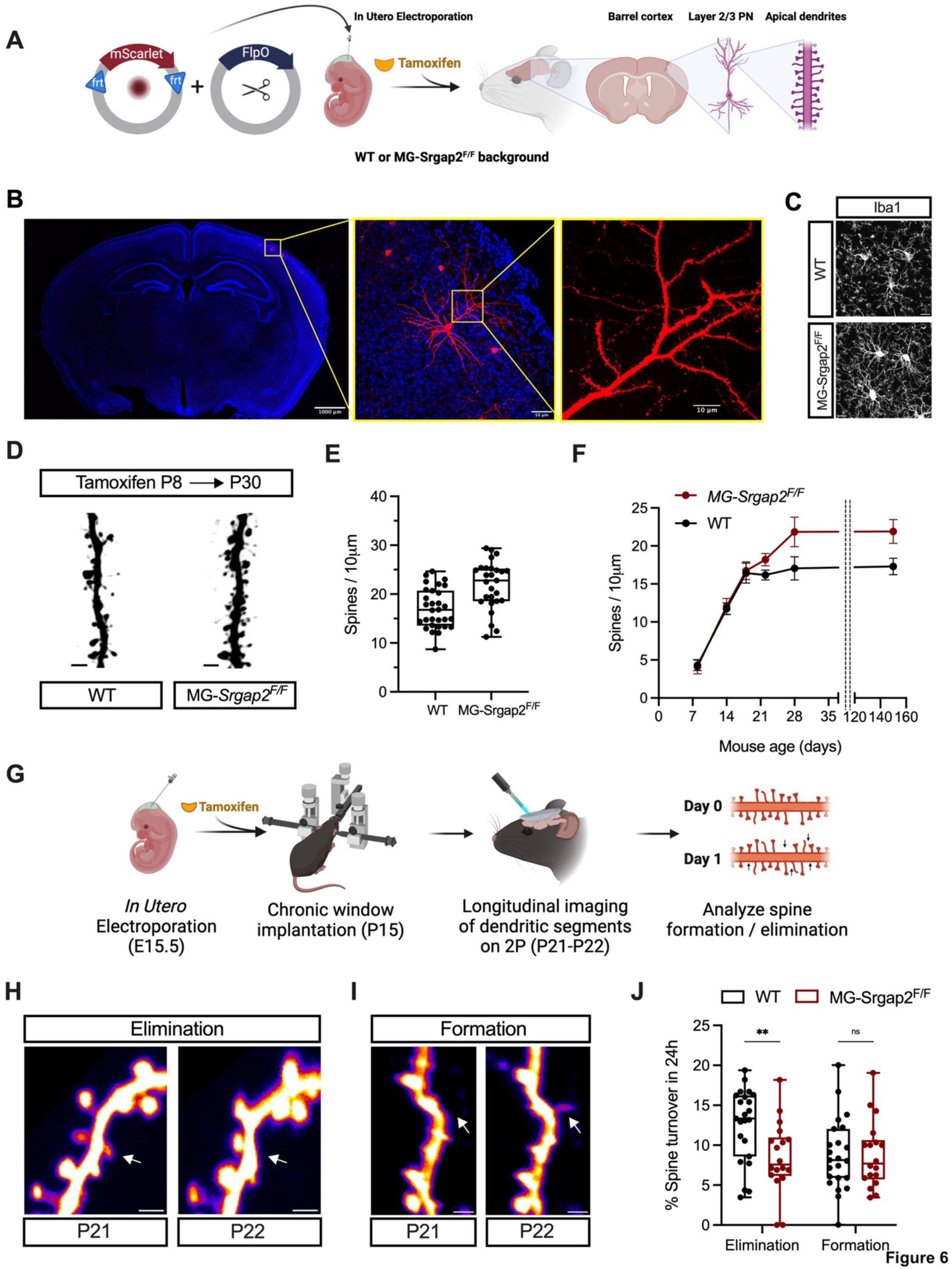
Microglial *Srgap2* regulates dendritic spine density and turnover in cortical neurons. (A) schematic representation of the experimental approach to obtain sparse fluorescent labeling of layer 2/3 pyramidal neurons (L2/3 PNs). (B) representative confocal images of a brain section containing sparsely labeled, mScarlet-expressing L2/3 PNs, and detail of its apical oblique dendrites featuring synaptic spines. (C) Representative images of Iba1^+^ microglia from P30 WT and *MG-Srgap2^F/F^* mice having undergone *in utero* Electroporation. D) Representative image detail of L2/3 PNs dendritic segments from P30 WT and *MG-Srgap2^F/F^* mice. Scale bar, 2 μm. E) Dendritic spine density from L2/3 PNs from these mice. Data from 2 dendritic segments per neuron, total of 14-16 neurons from 4-5 mice per genotype. (F) Dendritic spine density of L2/3 PNs from WT and *MG-Srgap2^F/F^* mice throughout mouse cortical development. Data from 7-22 neurons from 3-7 mice per genotype and timepoint. (G) Strategy to longitudinally image L2/3 PN dendritic segments *in vivo* in juvenile mice. (H-I) Representative examples of dendritic spine elimination (H) and formation (I) in L2/3 PN during a 24h time period. Scale bar, 2 μm. (J) Rates of dendritic spine formation and elimination in L2/3 PN from WT and *MG-Srgap2^F/F^* mice. Data from 3 mice per genotype, 18-23 segments per genotype.

Analysis of L2/3 PN apical oblique dendritic spines in *MG-Srgap2^F/F^* mice revealed a higher spine density at P30 compared to littermate controls (**Fig. 6D-E**). We then assessed when this increased spine density emerged during development, since P30 is considered a mature stage for synaptic development in WT conditions. In L2/3 PNs of WT mice, synaptic density quickly increased in the first 2 weeks after birth and remained stable afterwards, while in *MG-Srgap2^F/F^*neurons synaptic density increased monotonically during the first 4 weeks after birth to reach significantly higher density that is maintained in adult mice (**Fig. 6F**). This result suggests that induction of SRGAP2-dependent neotenic microglial maturation results in a cell non-autonomous, prolonged period of synaptic development in cortical pyramidal neurons, ultimately leading to higher synaptic density.

To determine if the increased dendritic spine density reached in mice carrying microglia-specific SRGAP2 loss-of-function results from increased spine formation and/or reduced spine elimination during the peak of synaptogenesis in L2/3 PNs, we performed longitudinal *in vivo* two-photon microscopy (**Fig. 6G**). We implanted a cranial window above the primary somatosensory cortex (S1) of juvenile (P15) wild-type control mice or littermate *MG-Srgap2^F/F^* mice induced with tamoxifen at P8-9, and tracked dendritic spine formation/elimination events in the apical dendrites of mScarlet-expressing L2/3 PN between P21 and P22 (**Fig. 6H-I**). We found that while both WT and in *MG-Srgap2^F/F^* neurons exhibited similar rates of spine formation, *MG-Srgap2^F/F^* neurons displayed lower rates of spine elimination (**Fig. 6J**), suggesting that the higher synaptic density observed in L2/3 PNs following microglia-specific Srgap2 deletion is due primarily to reduced synapse elimination.

## DISCUSSION

Altogether, our data indicates that *Srgap2* is essential for the terminal maturation of mouse and human microglia, and that delayed microglial maturation can lead to extended periods of synaptic development. In humans, the slow acquisition of a terminal maturation transcriptional signature and the appearance of *Srgap2* paralogs contribute to the neotenic development of microglia.

Recent scRNAseq studies have highlighted the developmental trajectories of human microglia during fetal and postnatal stages, although maturational trajectories over early postnatal development, and at longer timescales, are still not well understood ^31–34,57^. Our data suggest that human microglia acquire features of terminal maturation much slower than mouse microglia, and identify the gene Srgap2 as a master regulator of maturation and function.

While we have identified the function of one candidate human-specific genetic modifier of microglia, i.e. the human-specific paralogs *Srgap2b/c*, many other potential human-specific modifiers could also have played a role during evolution in the acquisition of human-specific features of microglia. These include an epigenetic developmental program setting the slow timing of human neuronal maturation ^58^; human-specific base pair substitutions in regulatory regions otherwise ultra-conserved throughout vertebrate evolution (termed Human Accelerated Regions) that might impact spatio-temporal patterns of gene expression ^59,60^; the occurence of human-specific mRNA splicing events ^61^; slower protein catalytic rates ^62^; or the presence of human-specific polymorphisms in genes involved in microglial function ^36,63^. Indeed, polymorphisms observed in the human *ApoE* and *Trem2* gene loci have been shown to endow hiPSC xenotransplanted microglia with distinct transcriptional responses to amyloid beta plaques ^36^, suggesting that this is another potential mechanism by which microglia can exhibit human-specific structural and/or functional features. Future studies will need to assess the impact of these other factors in the acquisition of human-specific features in microglia.

The finding that human cortical excitatory neurons exhibit increased number of synapses per neuron has been hypothesized to be a key factor underlying our higher cognitive abilities ^4,30,64–66^. Recently, using human cortical pyramidal neuron xenotransplantation and mouse genetic models, multiple groups have shown that their cell-intrinsic slow maturation rate is partly responsible for such increase, allowing synapses to form during an extended period of time ^27,30,50^. Indeed, previous work from our lab and others demonstrated that neuronal *Srgap2* cell-cell-autonomously regulates synapse density and the timing of synaptic maturation in mouse and human L2/3 PNs, and that induction of human-specific SRGAP2C expression in mouse L2/3 PNs leads to increased cortico-cortical connectivity, more reliable sensory-evoked responses, and improved behavioral performance in a sensory-discrimination task ^44–46^. Given the prominent role of microglia in fine-tuning synaptic connectivity during mouse postnatal brain development, it is likely that the same mechanisms are at play during human postnatal brain development, and our results argues that in the human cortex, the expression of human-specific genes *SRGAP2B/C* in microglia participates to the prolonged period of synaptic maturation and the increased synaptic density observed in human cortical neurons.

Our results highlight the fact that the timing of synaptic and microglial maturation must be synchronized in a species-specific manner during brain development. Indeed, our data suggest that the temporally-restricted functions of microglia during mouse brain development are developmentally matched between mouse and human –albeit at strikingly different absolute timescales– and our estimated timing for human microglia maturation largely coincides with the reported timing of synapse maturation during human brain development ^4,6,15,41^.

Our results also highlight a set of genes (module 160) characterized by a temporal pattern of expression coinciding with predicted maturation of human microglia. Interestingly, these genes are overrepresented in transcriptomic signatures from primary human microglia coming from patients with Alzheimer’s disease ^67^. Given the specificity of this gene signature for primary human microglia ^36^, it is possible that their impact of these genes on microglia functional and structural maturation has been previously overlooked. Future work will shed light on the role of the genes in this Module 160 in shaping the human microglia response to neurodegenerative diseases.

### Limitations of the study

Compared to other hiPSC-derived microglia models (such as *in vitro* or organoid-based co-cultures) the hMG xenostransplantation model is considered to mirror most closely the postnatal development of human microglia, both morphologically and transcriptomically ^31,39,40,68^. However, hMGs still reside and develop within a mouse brain environment, and as such, the role of extracellular cues on microglia development may also depend on species-specific interactions. Mainly, the impact of the ectopically provided human Csf-1 on microglial development has not been carefully evaluated and may preclude the generalization of some of our findings. In this sense, a new xenotransplantable, tri-culture organoid system was recently developed where most of the extracellular cues (such as Csf-1) can be provided by its human astrocyte and neuron environment ^69^. Interestingly, mouse microglia development has been shown to be partially dependent on cues from the microbiota and their interaction with other immune cells, mainly CD4^+^ T cells ^70,71^. Whether such interactions play a role in human microglial maturation, and whether they are not fully captured in the hMG xenotransplantation model remains to be addressed. The relatively short lifespan of a laboratory mouse (∼2 years) currently limits our ability to fully assess hMG maturation longitudinally and its relation with human primary microglia, as exemplified by our inability to test whether hMGs ever reach more simplified morphology even ten months after xenotransplantation.

Finally, our analyses have been limited to microglia from the primary somatosensory cortex, given the role of this region in high-level associative and cognitive functions. Although it is widely accepted that fully mature, homeostatic microglia do not exhibit a high range of transcriptomic heterogeneity ^34,57,72^, it will be important to test whether microglia maturation respond differently to their environment other brain areas ^73^. For example, it has been shown that cerebellar microglia exhibit site-specific functional and developmental differences compared to cortical microglia ^74,75^. Future research will elucidate whether the molecular mechanisms described in our study play a role in other brain regions.

## METHODS

### Mice

All animals were handled according to protocols approved by the institutional animal care and use committee (IACUC) at Columbia University, New York. All experiments involving xenotransplantations were performed according to the standards set forth by the Department of Comparative Medicine and Massachusetts Institute of Technology. All mice used in these experiments were maintained on a regular 12 h light/dark cycle at 20 to 22 °C, humidity between 30 to 70%, and given feed and water *ad libitum*. Tmem119^creERT2^ mice ^52^ and *Ai9* mice ^53^ were obtained from Jax. Recipient mice for xenotransplantation (*Rag2^-/-^IL2rg^-/-^* mice carrying the transgene for the human Csf1 allele, C;129S4-Rag2tm1.1Flv Csf1tm1(CSF1) Flv Il2rgtm1.1Flv/J) were obtained from Jackson Labs. *Srgap2^-/-^*^44^, *Srgap2^fl/fl^* ^51^, and *Srgap2c^lsl^* ^46^ mice were generated in house. All mice used to compare differences across genotypes were littermates (except those generated to analyze microglial surveillance) and received the same tamoxifen dosage, and genotype was blinded to the experimenter. Individual strains were kept in a C57BL/6J background crossed with the outbred strain 129S2/SvPasCrl mice, for the purpose of improving pup viability after *in utero* electroporation. For experiments performed on adult mice (>P28) mice received two doses (75 μg/g body weight, BW) of tamoxifen (Sigma-Aldrich) at P7-8. For experiments performed on younger mice (<P28), mice received two doses (50 μg/g BW) of 4-hydroxytamoxifen (TargetMol) at P2-3. hMG and Sgrap2^-/-^ mice did not receive any tamoxifen.

### *In utero* Electroporation

*In Utero* Electroporation (IUE) was performed on isofluorane-anesthetized timed-pregnant females at E15.5 as previously described ^76^. Briefly, after exposing the embryos from the abdominal cavity, an endotoxin-free DNA mix containing dose-limiting amounts (5 ng/μl for Figures 7B-E, and 20 ng/μl for Figures 7F-I) of a FlpO plasmid, and 1 μg/μl of a flp-dependent fluorescent protein-encoding plasmid (mScarlet for Figures 7B-E, and mGreenLantern for Figures 7F-I) were injected into the ventricles of mouse embryos using a heat-pulled glass pipette. mScarlet-FRT and mGreenLantern-FRT plasmids were generated by subcloning the mScarlet and mGreenLantern coding sequence (Addgene #85042 and #164468, respectively) inside the FRT-flanking region of a CAG-promoter plasmid (Addgene #99133). Electroporation into cortical neurons was performed applying 4 pulses of 42 V for 50 ms with 500 ms intervals, using a 3 mm diameter platinum tweezer electrode (Nepa Gene) and a square wave electroporator (ECM 830, BTX). Embryos were placed back into the abdominal cavity and the incision was sutured, allowing the mouse to recover on a heated pad and give birth naturally.

### Cortical Window Implantation

Adult (∼P50 for Figures5A-C) or juvenile (P15, for figures 7F-I) mice were anesthetized with isofluorane and given analgesic (rimadyl, 10 μg/g BW) and topical anesthetic (lidocaine 10 μg/g BW), prior to exposing the dorsal skull with a scissors. After performing a craniotomy, a glass window (3 mm diameter, Warner Instruments) was placed over the primary somatosensory cortex and secured with cyanoacrylate (Loctite super glue). Ice-cold sterile artificial cerebrospinal fluid (aCSF, Tocris) was applied when necessary to prevent excessive heating and dryness throughout the craniotomy. A custom 3D-printed headplate was secured onto the skull using dental adhesive cement (Parkell). Coverslip was protected with silicone (World Precision Instruments) until imaging, and reapplied between imaging sessions. After surgery, mice were injected with 20 μl/g BW of sterile PBS and allowed to recover under a heated pad. Mice received two additional daily analgesic injections after surgery and were imaged after 2 weeks for adult mice, and 6 days for juvenile mice.

### Lentivirus production

Vpx-containing Lentiviruses were produced as described ^55^. Briefly, HEK293 cells were cultured in DMEM High Glucose (Cytiva) according to standard protocol and transfected using polyethylenimine (PEI) with the following plasmids: pMDL-X, VSV-G, HIVRev, Vpx, and shRNA-encoding plasmid at a mass ratio of 10:7:5:2:28. pMDL-X, VSV-G, HIVRev, Vpx –encoding plasmids were a gift of Dr. Nathaniel Landau. Plasmids encoding shRNA against SRGAP2B/C and scramble shRNA were generated and previously validated ^44,50^. 72 hours after transfection, the viral supernatant was collected, centrifuged at 2,000rpm for 5 min to remove cell debris, filtered through a 0.45 μm PES membrane, and further concentrated by centrifuging at 16,000 rpm for 2h at 4°C on a tabletop centrifuge. Pellets were stored at –80°C. Viral titers were estimated using Lenti-X GoStix tests (Takara Bio).

### Human induced pluripotent stem cell (hiPSC) culture, myeloid precursor differentiation, lentiviral infection and xenotransplantation

HiPSCs lines used in this study were previously generated and characterized in the Jaenisch Lab ^77^. The cell lines used for the morphological characterization of wild-type human microglia, WIBR-hiPS-SNCA ^A53T-Corr^, stably expressed Green Fluorescent Protein (GFP) from the AAVS1-locus after targeting with a Zinc-Finger GFP-expressing AAV, as previously described ^78,79^. The cell line used for *Srgap2b/c* knock-down experiments, WIBR-hiPS-SNCA ^A53T-Corr^, was unlabeled. HiPSCs lines were maintained in feeder-free conditions and regularly tested for mycoplasma contamination. HiPSCs were differentiated into myeloid precursors (MPs) using a previously published protocol ^79,80^. Briefly, HiPSCs were cultured for four days in embryoid body medium (containing ROCK inhibitor, BMP-4, SCF, and VEGF-121), and then cultured in hematopoietic medium (containing M-CSF and IL-3) for a maximum of 1.5 months, during which cells were harvested. For *Srgap2b/c* knock-down experiments, cultured MPs were infected a mix of lentivirus and protamine sulfate (5 µg/mL, Thermo Fisher Scientific) to enhance lentiviral integration into the cells. and infected cells were subsequently selected with zeocin (Thermo Fisher scientific) at 1 mg/mL 72 hours after infection. For xenotransplantation, cultured MPs were resuspended in phosphate-buffered saline without calcium and magnesium prior to injection at a concentration of 10^5^ cells/µL. Postnatal day 0 (P0) to P3 mouse pups of either sex were manually injected with 4.10^5^ MPs in the lateral ventricles (1 anterior and 1 posterior injection site per brain hemisphere) using glass micropipettes.

### Slice preparation, Immunostaining, confocal imaging and analysis

At the indicated age, mice were anesthetized with isofluorane and perfused intracardially with 4% paraformaldehyde (PFA) (Electron Microscopy Sciences). Brain was left to fix on PFA for 2h at room temperature (RT) and then wash on PBS at 4°C over night (O/N). Brains were sectioned coronally at 100um thickness on a vibratome 9Leica VT1200S). When immunostained, slices were incubated in blocking buffer (5% Goat Serum, 0.5% Triton X in PBS) for 30 min at RT, and stained with their respective primary antibodies (Iba1 clone E4O4W, Cell Signaling; and CD68 Clone FA-11, BioRad) in staining buffer (2% Goat Serum, 0.5% Triton X in PBS) at RT O/N. The following day, slices were washed twice with PBS and incubated with fluorescent secondary antibody in staining buffer. Slices were finally washed and incubated in DAPI (1:5000 in PBS) and mounted on slides with Prolong Glass Antifade (Thermo Fisher Scientific). All antibodies used in this study are listed on the reagent’s table. Slices for microglia morphological analysis were imaged using a Nikon A1 laser-scanning confocal microscope controlled by NIS-elements (Nikon) at 1024×1024 using a 20x Plan Apo 0.8 NA objective (Nikon) to quantify microglia numbers and a 60x Plan Apo 1.4 NA oil immersion objective (Nikon) to quantify their morphology. For morphology measurements, a 12 μm z-section was acquired with a 0.3 μm z-step, and later MaxIP for analysis. For experiments involving dendritic spine quantification, images were acquired on a CSU-W1 Yokogawa Spinning Disk Field Scanning Confocal System (Nikon) using a 100x Plan Apo 1.35 NA silicone-immersion objective (Nikon). Images were acquired with a 0.15 μm z-step and variable z-section thickness dependent on dendritic segment orientation. Microglial morphological complexity from MaxIP images was quantified on Fiji ImageJ using 2D sholl analysis. Briefly, individual microglia were manually cropped, binarized while excluding particles <20pixels, and a pointer was added to the cell center for sholl analysis. Dendritic spine density from L2/3 pyramidal neurons was manually quantified across the z-section (3D). Large brain slice-wide images were obtained with a 4X Plan Apo 0.2 NA (Nikon).

### *In vivo* 2-photon microscopy: image acquisition, processing, and analysis

Animals were imaged under isofluorane anesthesia on a custom-built Bergamo II two-photon microscope running ThorImage LS ver. 4.1 and Thorsync (Thorlabs), using a 25x SCALEVIEW-A2 1.0 NA objective (Olympus) at 920 nm wavelength (for imaging mGreenLantern, Figures 7F-I) or 980 nm wavelength (for imaging Tdtomato, Figure 5A-C) using a Ti-Sapphire laser (Coherent). Imaging was performed ∼30-100 μm below the pial surface at 1 μm z-step. For microglia surveillance (Figures 5A-C), images (20 cumulative frames) were acquired at 3X zoom with 0.155 μm/pixel XY resolution, imaging a 20 μm z-section each minute for 10 minutes. For dendritic spine longitudinal imaging (Figures 7F-I), regions of interest were manually re-located 24h after initial imaging based on overlaying cranial blood vessel architecture, and images (50 cumulative frames) were acquired at 4.1X zoom with 0.117 μm/pixel XY resolution and variable z-section thickness dependent on dendritic segment orientation. Motion correction and Image registration were performed on Fiji ImageJ. Still images (Figures 7F-I) were aligned using rigid body registration with the StackReg Plugin. Time-lapse images (Fig 5A-C) were converted to Hyperstacks and aligned using rigid body registration with the HyperStackReg Plugin. Registered images were bleach-corrected using histogram-matching and converted to Maximum Intensity Projections (MaxIP) of each z-stack. Using the MaxIP time-lapse images, Microglia surveillance was then quantified using the MotiQ 2D Analyzer Plugin ^54^. Dendritic spines were quantified manually across the z-section (3D), and individual spines were matched across days. Spine dynamics were quantified as percentage of spines eliminated or formed on day 2 compared to day 1.

### WGCNA and K-means clustering

#### Module generation and analysis using WGCNA

Transcriptional signatures of human and mouse microglia were obtained using human datasets from ^41^ (available at https://pre-postnatal-cortex.cells.ucsc.edu) and mouse datasets from ^42^ (GEO series GSE200202) and generating subsets containing exclusively microglia. The human microglia subsets were generated from ^41^ by binning the multiple development stages in the original dataset into ‘stage’ groups to reflect developmental milestones and to include comparable number of microglia per group. The mouse microglia subsets were generated from ^42^ maintaining the annotated developmental stages but including exclusively cells annotated as “Juvenile” or “Adult” microglia. Modules were generated from datasets using WGCNA (ver. 1.72.5 on R 4.3.1) as previously described ^81^. The activity of these gene modules was evaluated using the module activity metric described in ^82^, in which the activity of each module is evaluated for each cell by calculating the average normalized counts per million (CPM) detected for each module gene.

#### Grouping WGCNA module by K-means clustering

The WGCNA module activity value for each module was used to perform k-means clustering (packages factoextra_1.0.7, cluster_2.1.6 and dplyr_1.1.4 on R 4.3.2) to assign the modules to different groups. The number of clusters were determined by calculating within-cluster squared errors (WSS) and the elbow methods, to choose a number of clusters whereby adding another cluster does not further improve the total WSS.

### Statistical analysis

All quantitative data with a single timepoint is represented as a boxplot, with whiskers indicating the maximum and minimum value, a horizontal line indicating the median, and all datapoints displayed. All temporal (except from transcriptomic data) and Sholl curve data shows the group mean of each timepoint and genotype, and error bars indicate the 95% confidence interval. Statistical analyses were performed using standard t-test when only two groups were compared, one-way ANOVA when three or more conditions were compared across the same magnitude, or two-way ANOVA when two conditions were compared across two magnitudes. Statistical significance was represented according to GraphPad format: ns, p> 0.05; *, p≤ 0.05; **, p≤ 0.01; ***, p≤ 0.001; ****, p≤ 0.0001. Data was plotted and analyzed using Prism software except for K-means clustering and transcriptomic data from Fig S2.

## ACKNOWLEDGEMENTS

We thank Sergio Bernal Garcia for valuable input and technical assistance during optimization of image acquisition and processing of 2-photon microscopy data. We thank Cynthia Jia for support during image data analysis. We thank Prof. Dorothy Schafer for insightful discussions and experimental guidance. We thank Martina Proietti Onori, Sergio Bernal Garcia, Aleksandra Recupero, and other Polleux lab members for extensive discussions. Schematic graphics were generated with BioRender. We thank Qiaolian Liu for excellent technical assistance with mouse colony management. Imaging was performed in the Zuckerman Institute’s Cellular Imaging platform. This work is supported by a Revson Senior Fellowship in Biomedical Sciences (CDS), a career award from the NIH-NINDS (R35 NS127232) (FP), and an award from the NOMIS Foundation (FP). This work was also supported by an award from the NIH-NIMH (R01 MH104610) (RJ), NIH-NINDS (R00 NS111731) (AB) and a Broad Stem Cell Research Center Fellowship (PN).

## AUTHOR CONTRIBUTIONS

Conceptualization, CDS, FP; Methodology, CDS, MK, JY, PN, FP; Human iPSC work and xenotransplantation, MK; scRNAseq data analysis, JY, PN; Investigation, CDS, JY, PN; Resources, CDS, FP, AB, RJ. Project administration, CDS, FP; Supervision, FP, AB; Funding acquisition, CDS, MK, RJ, FP. Writing – original draft: CDS, JY, PN, FP.

---

KEY RESOURCES TABLE.

KEY RESOURCES TABLE
| REAGENT OR RESOURCE | SOURCE | IDENTIFIER |
| --- | --- | --- |
| **Antibodies** |  |  |
| Iba1 (clone E4O4W) | Cell Signaling | 17198S |
| CD68 (Clone FA-11) | BioRad | MCA1957 |
| CD45 (clone 104) | Tonbo | 25-0454 |
| CD11b (clone M1/70) | Tonbo | 20-0112 |
| CX3CR1 (clone SA011F11) | Biolegend | 149015 |
| Alexa488 Goat $\alpha$ Rabbit | Thermo Fisher Scientific | A-11008 |
| Alexa555 Goat $\alpha$ Rat | Thermo Fisher Scientific | A-21434 |
| Alexa555 Goat $\alpha$ Rabbit | Thermo Fisher Scientific | A-21428 |
| CF568 Goat $\alpha$ Rat | Biotium | 20096 |
| CF568 Goat $\alpha$ Rabbit | Biotium | 20801 |
| **Chemicals, Recombinant proteins, consumables, etc.** |  |  |
| Phosphate Buffer Saline (PBS) | Cytiva | SH30256.FS |
| Goat Serum | Invitrogen | 16210-064 |
| Fetal Bovine Serum | Thermo Fisher Scientific | 16141002 |
| Triton X-100 | Sigma-Aldrich | T9284 |
| 4',6 Diamidino 2 Phenylindole, Dihydrochloride (DAPI) | Thermo Fisher Scientific | D1306 |
| Formaldehyde Aqueous Solution, 32%, Liquid (PFA) | Electron Microscopy Solutions | 15714-S |
| Prolong Glass Antifade | Thermo Fisher Scientific | P36980 |
| Tamoxifen | Sigma-Aldrich | T5648 |
| (Z)-4-hydroxy Tamoxifen | TargetMol | T4420 |
| TRIzol | Thermo Fisher Scientific | 15-596-018 |
| Percoll | Sigma-Aldrich | P4937 |
| NEBNext® Ultra™ II RNA Library Prep kit | New England Biolabs | E7770S |
| NEBNext Poly(A) mRNA Magnetic Isolation Module | New England Biolabs | E7490 |
| NEBNex Multiplex Oligos for Illumina | New England Biolabs | E7335S |
| ROCK inhibitor | EMD Millipore | 688000 |
| BMP-4 | Thermo Fisher Scientific | RHBMP-4 |
| SCF | Life Technologies | PHC2115 |
| VEGF-121 | Life Technologies | PHC9394 |
| M-CSF | Life Technologies | PHC2015 |
| IL-3 | Life Technologies | PHC0034 |
| Protamine sulfate | Thermo Scientific | J62926.06 |
| Zeocin | Thermo Scientific | R25001 |
| Dulbecco's Modified Eagle Medium (DMEM) | Cytiva | 05890 |
| Lenti-X GoStix Plus tests | Takara Bio | 631280 |
| Polyethylenimine, 25000 Molecular Weight, Linear (PEI) | Polysciences | 23966-100 |
| Low toxicity silicone adhesive | World Precision Instruments | KWIK-SIL |
| Parkell C&B Metabond Quick Adhesive Cement System (Dental cement) | Parkell | 04-PKL-S380 |
| Artificial Cerebrospinal Fluid (aCSF) | Tocris | 3525 |
| 3 mm round coverglasss, #0 thickness | Warner Instruments | 64-0726 |
| Loctite Super Glue Ultra Liquid control | Loctite | 1647358 |
| **PLASMIDS and cell lines** |  |  |
| HEK-293T cell line | ATCC | CRL-3216 |
| WIBR-hiPS-SNCA<sup>A53T-Corr</sup> | Jaenisch Lab<sup>77</sup> | N/A |
| pAAV-CAGGS-EGFP | Addgene | #22212 |
| pCAG-FlpO | Addgene | #60662 |
| pAAV-CAG-fDIO-mNeonGreen | Addgene | #99133 |
| pAAV-EF1a-DIO-mGreenLantern | Addgene | #164468 |
| pmScarlet_C1 | Addgene | #85042 |
| pAAV-CAG-fDIO-mGreenLantern-FRT (mGreenLantern-FRT) | This study | N/A |
| pAAV-CAG-fDIO-mScarlet-FRT (mScarlet-FRT) | This study | N/A |
| Sh RNA Scramble | Polleux Lab<sup>44</sup> | N/A |
| Sh RNA targeting SRGAP2B/C | Polleux Lab<sup>44</sup> | N/A |
| pMDL-X | Landau Lab<sup>55</sup> | N/A |
| VSV-G | Landau Lab<sup>55</sup> | N/A |
| HIVRev | Landau Lab<sup>55</sup> | N/A |
| Vpx | Landau Lab<sup>55</sup> | N/A |

| REAGENT OR RESOURCE | SOURCE | IDENTIFIER |
| --- | --- | --- |

| MOUSE STRAINS |  |  |
| --- | --- | --- |
| 129S2/SvPasCrl mice (129 mice) | Charles River | 287 |
| B6.Cg-Gt(*ROSA*)26Sor<sup>tm9(CAG-tdTomato)Hze/J</sup> (Ai9) | The Jackson Laboratory | 007909 |
| C57BL/6-*Tmem119*<sup>em1(*cre/ERT2*)Gfng/J</sup><br>(*Tmem119*<sup>creERT2</sup>) | The Jackson Laboratory | 031820 |
| Constitutive *Srgap2* knockout mouse | Polleux Lab <sup>44</sup> | N/A |
| Conditional *SRGAP2* knockout mouse<br>(*Srgap2*<sup>F/F</sup>) | Polleux Lab <sup>51</sup> | N/A |
| *Rosa26*<sup>SRGAP2C(F/+)</sup> knockin mouse (*Srgap2c*<sup>LSL</sup>) | Polleux Lab <sup>46</sup> | N/A |
| C;129S4-*Rag2*<sup>tm1.1Flv</sup> *Csf1*<sup>tm1(CSF1)Flv</sup> *Il2rg*<sup>tm1.1Flv/J</sup><br>(*Rag2*<sup>-/-</sup> *il2rg*<sup>-/-</sup> hCsf-1 knock-in) | The Jackson Laboratory | 017708 |

## Supplementary Figures

Diaz-Salazar et al. (2024)

Four Supplementary figures (S1-4)

**Figure S1.**
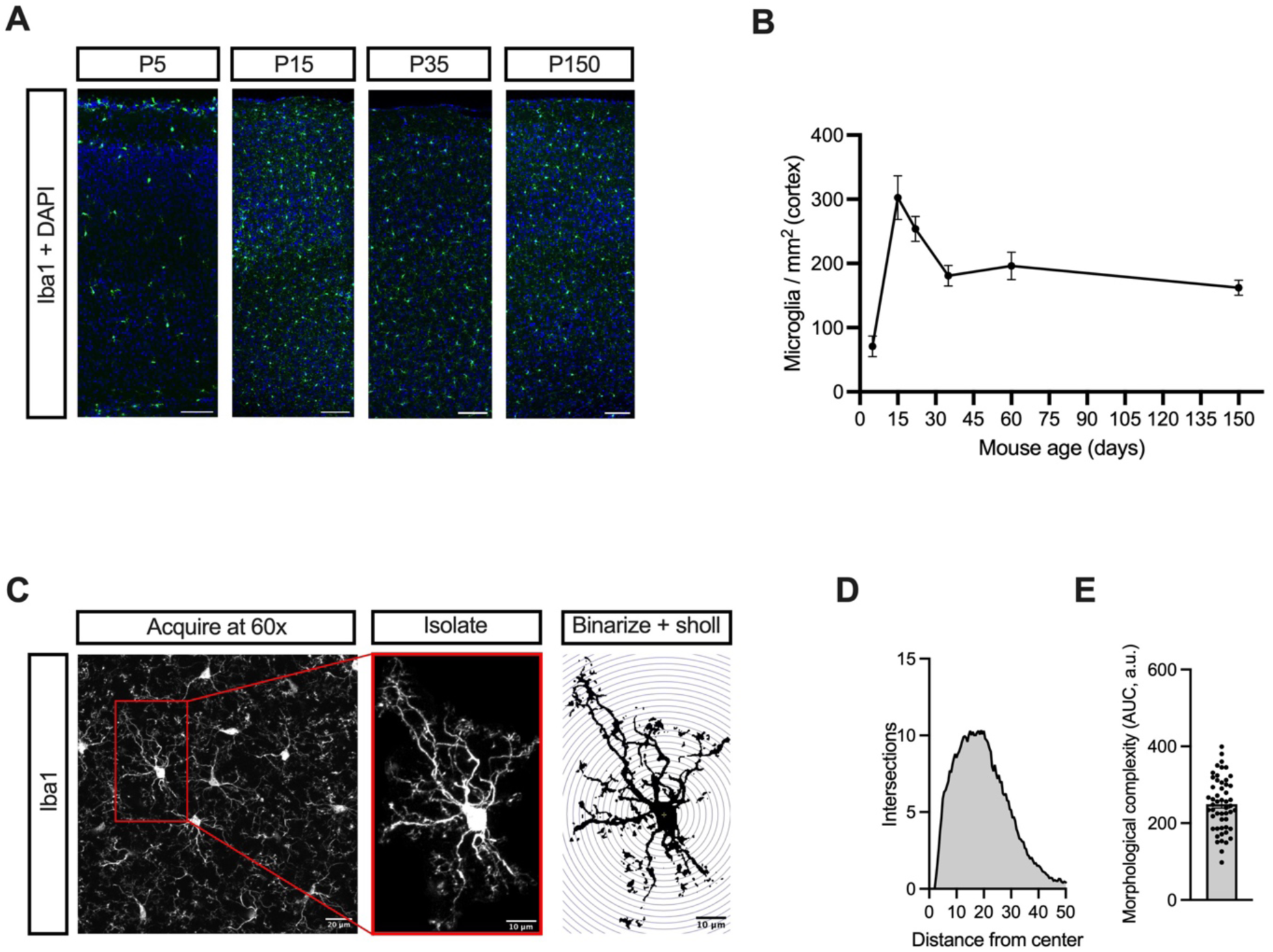
Approach to analyze microglial complexity during development (related to Figure 1). (A) Representative images of cortical microglial density in neonatal (P5), juvenile (P15), early adult (P35), and late adult (P150) mice. Scale bar, 100 μm. (B) Quantification of cortical microglial density throughout development. Data from 3 to 7 mice per timepoint, 2 to 4 fields per mouse. (C-E) Analytical approach to assess microglial morphology. Microglia (as revealed by Iba1 staining) from layer 2/3 of the barrel cortex were imaged using confocal microscopy (C), isolated, binarized, and subjected to Sholl analysis (C), counting the intersections of concentric circles with microglia branches (D). The Area under the curve (AUC) of the Sholl traces was computed as a measure of microglial morphological complexity (E).

**Figure S2.**
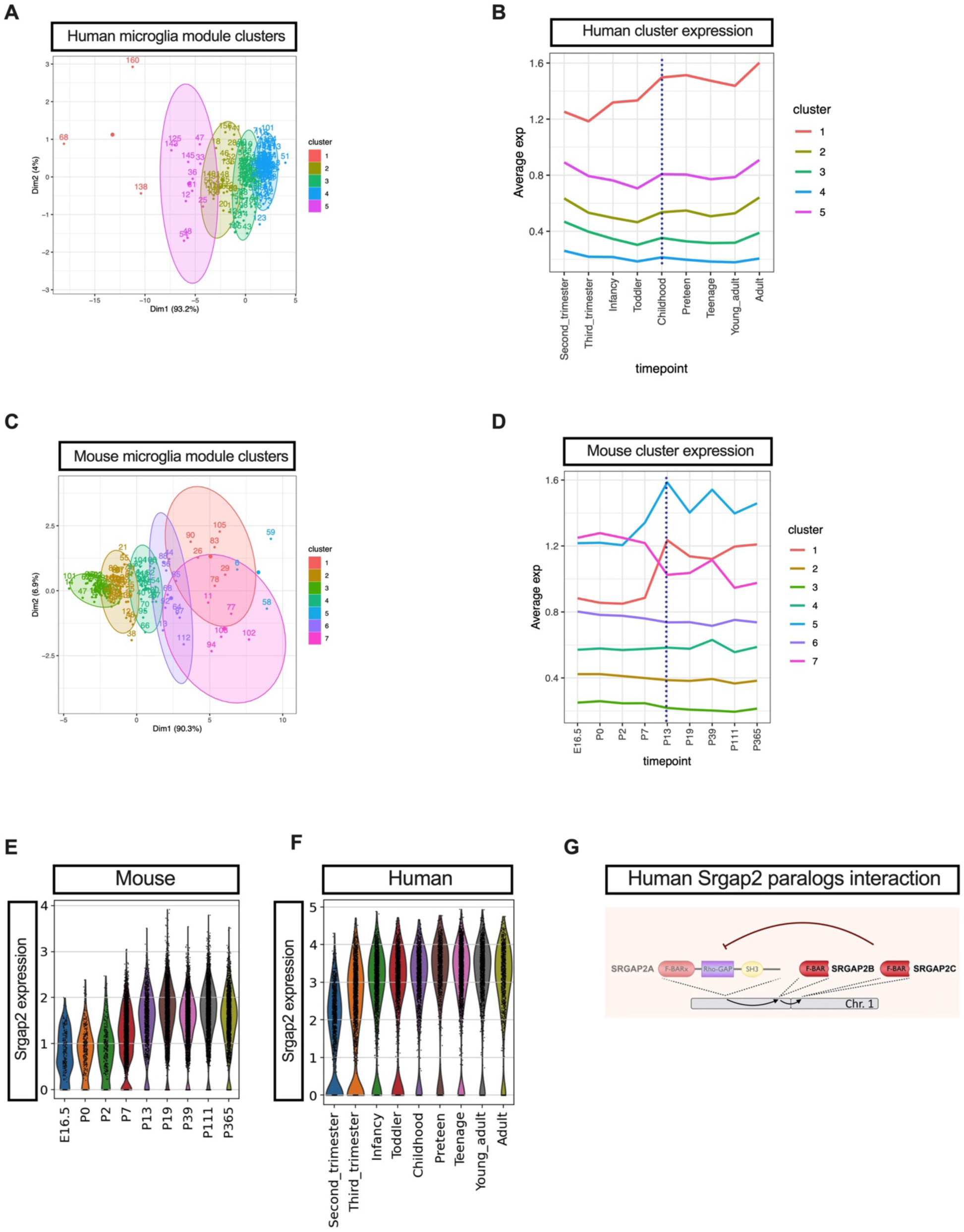
Unsupervised analysis of human and mouse microglial transcriptomic dynamics during development and maturation (related to Figure 2). (A and C) k-Means clustering of Weighted Gene Correlation Network Analysis (WGCNA) modules of co-expressed genes during human (A) or mouse (C) microglia development. (B and D) Average activity of module clusters throughout human (B) or mouse (D) microglia development. (E-F) Violin plot depicting expression of the *Srgap2* gene in mouse (E) and human (F) microglia throughout development. (G) Scheme of Srgap2 partial gene duplications and protein interactions, depicting partial inhibition of Srgap2 functions by its human-specific paralogs.

**Figure S3.**
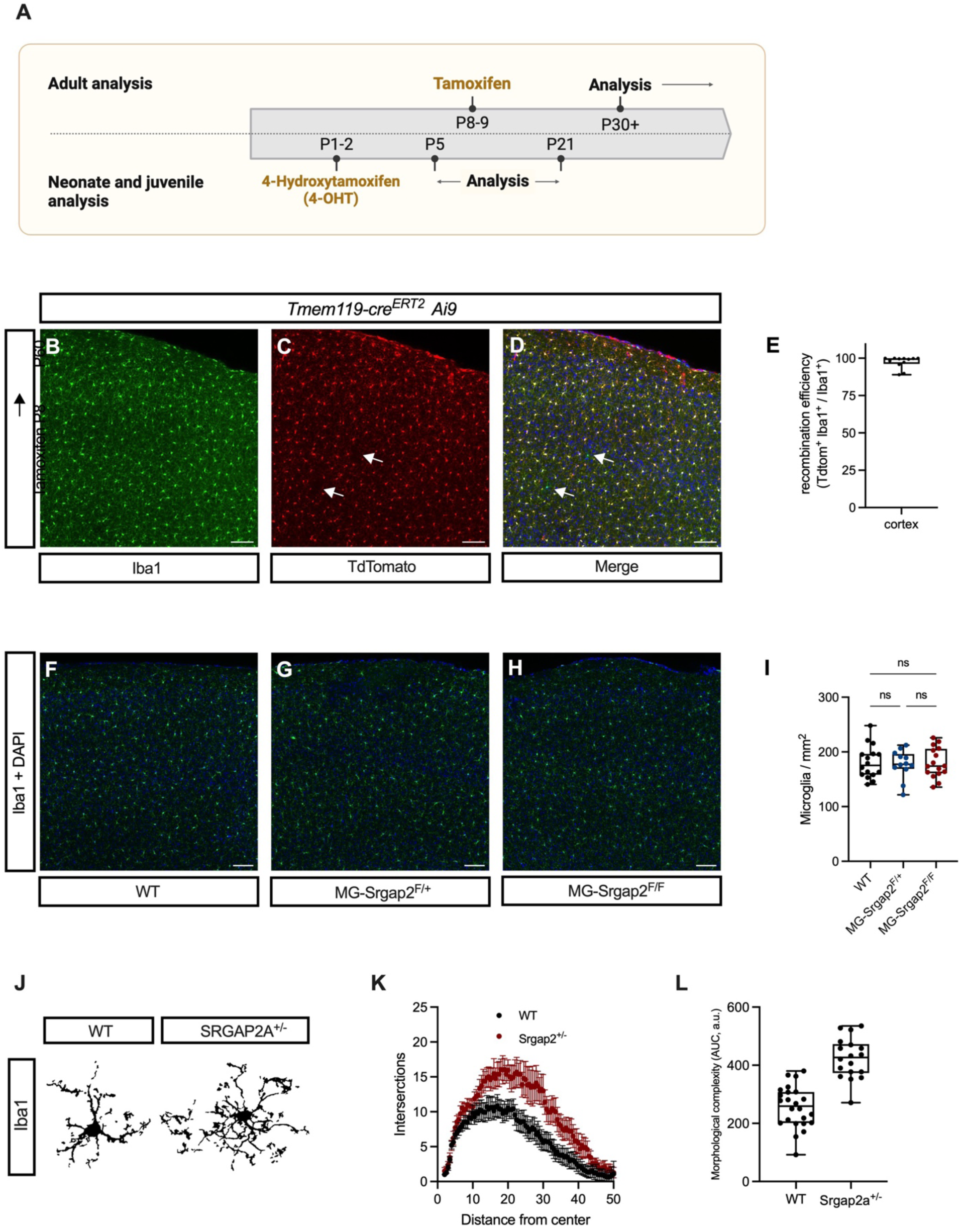
Microglial density, Tamoxifen dosage, and morphological complexity in constitutive knockout Srgap2+/− mice (related to Figure 3). (A) Tamoxifen dosing approach employed to drive cre-mediated srgap2 gene excision in mouse microglia, for analysis at mature stages or during development. (B-D) Representative images of cortical microglia from P60 *Tmem119CreErt2 crossed with Ai9F/+* mice, showing high recombination efficiency upon tamoxifen administration at P8-9. White arrows, rare events of Iba1-positive but tdTomato-negative microglia where Cre-dependent recombination did not occur. Scale bar, 10 μm. (E) Quantification of recombination efficiency in microglia from these mice. Data from four mice, three fields per mouse. (F-I) Representative images of cortical microglial in adult WT, *MG-Srgap2F/+* and *MG-Srgap2F/F* mice. Scale bar, 100 μm. (J) Quantification of cortical microglia density in these mice. Data from 4 to 6 mice per genotype, 2 to 3 fields per mouse. (K) Representative binary traces of microglia from constitutive Srgap2+/− mice and littermate control WT mice. (L-M) Comparison of Sholl analysis (L) and morphological complexity (M) of microglia from these mice. Data from 2 mice per genotype, 8-14 cells per mouse.

**Figure S4.**
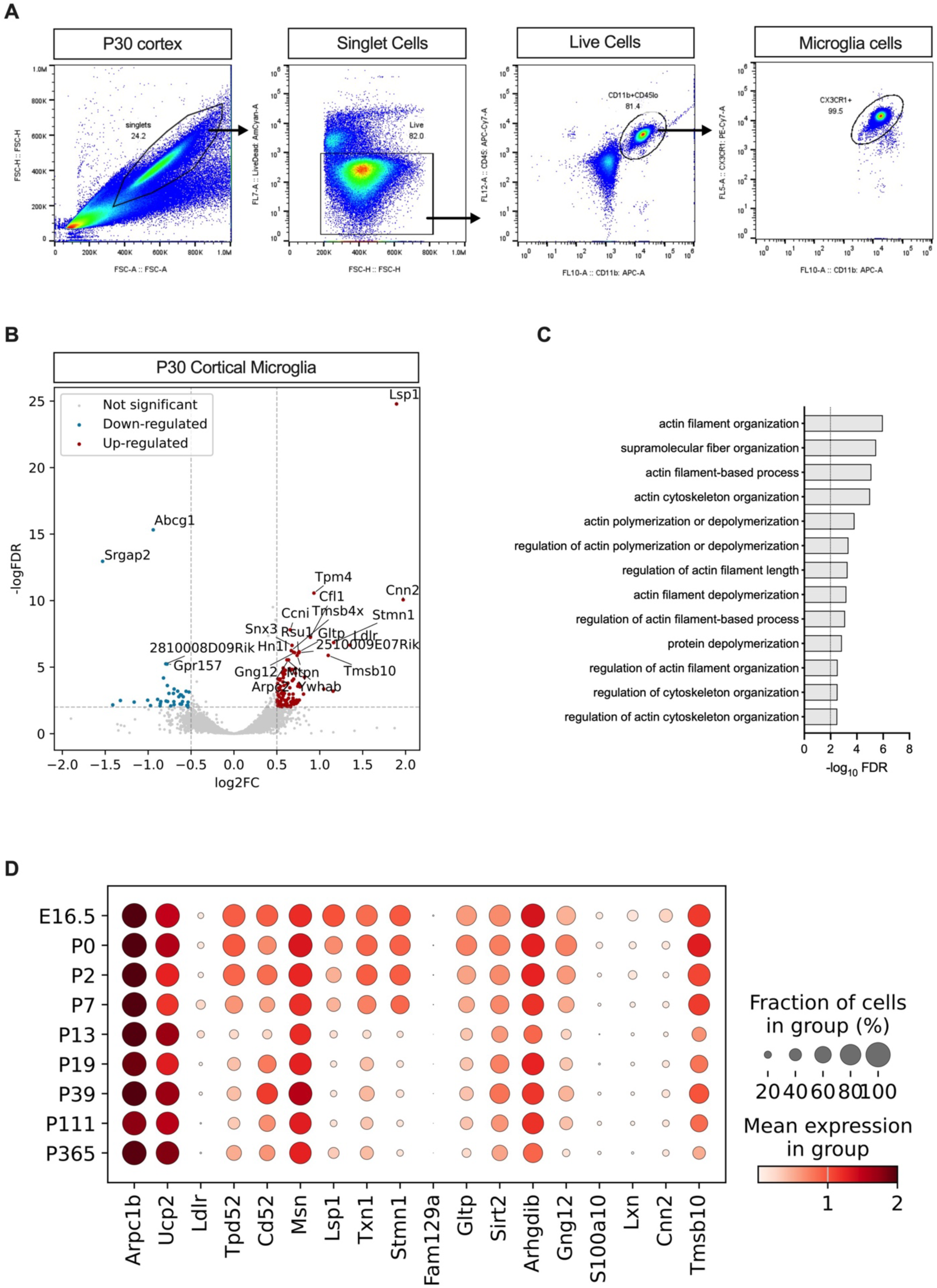
Srgap2 is required to acquire a mature transcriptomic signature in microglia (related to Figure 4). (A) FACS gating strategy to sort cortical microglia from P30 cortices for bulk RNA sequencing. (B) Volcano plot representing differentially expressed genes between WT and *MG-Srgap2^F/F^*microglia obtained by bulk RNA sequencing of FACS-sorted cortical microglia. Data from 3-4 biological replicates (individual mice). (C) Gene Ontology enrichment analysis displaying overrepresented pathways in *MG-Srgap2^F/^*^F^ microglia at P30, when compared to WT microglia. (D) Developmental expression profile of top overexpressed genes in *MG-Srgap2^F/^*^F^ microglia.

## EXTENDED METHODS

### RNA sequencing sample preparation, library generation, and data analysis

Fresh cortices from ice-cold PBS-perfused anesthetized mice were quickly homogenized made into a single-cell suspension by passing it through a 100 μm filter. Immune cells were isolated by centrifuging a brain homogenate on a 40% Percoll gradient at 2000 rpm for 5 min at 4°C. Myelin and fat debris were removed, cells washed with PBS, and incubated (2% FBS in PBS) with the following antibodies for 30 min at 4°C: CD11b-APC (1:200,Tonbo), CD45-APCCy7 (1:200 Tonbo), Cx3Cr1-PECy7 (1:200 Biolegend). After washing twice, microglia (∼100,000 cells / cortex) were sorted directly into TRIzol (Thermo Fisher Scientific) as CD11b^+^, CD45^low^, CX3CR1^+^ cells on a Sony MA900 (SONY) sorter with a 100 μm microfluidic chip. TRIzol containing the cell lysates was mixed with chloroform at a 5/1 ratio, shaken and spun at 12,000 g for 15 min at 4°C. Clear supernatant was transferred to a fresh tube and washed with 70% ethanol. RNA cleanup was performed on a RNeasy Mini (Qiagen) according to manufacturer instructions. RNA quantity and quality was assessed with a 2100 Bioanalyzer (Agilent). cDNA Libraries were prepared from samples with >8.5 RIN using the NEBNext Ultra II RNA Library Prep quit, the NEBNext Poly(A) mRNA Magnetic Isolation beads, and the NEBNex Multiplex Oligos for Illumina (all New England Biolabs), all according to manufacturer’s instructions. Samples were run on a NextSeq 2000 analyzer (illumina) according to manufacturer’s instructions. Reads were aligned to the GRCm39 mouse genome using BaseSpace DRAGEN RNA app (Illumina). Differential gene expression analysis was performed with pyDESeq2 ^81,82^. Genes with less than 10 read counts in total were filtered out. ERT2 expression from the Tmem119^creERT2^ locus was mapped to the Esr1 gene and was thus filtered out (log2FC=xx, padj=ppp) for visualization. Genes with |Log2FC (log_2_ fold change)| > 0.5 and adjusted p-value (logFDR) < 0.01 between MG-Srgap2^F/F^ and WT samples were assigned as up-or down-regulated.

